# Direct and indirect regulation of fetal globin transcript by RNA-binding protein IGF2BP1

**DOI:** 10.1101/2025.09.09.675143

**Authors:** Steven Coyne, Ting Wu, Mir Hossain, Divya Vinjamur, Marlena Starrs, Jing Zeng, Felicia Andresen, Ashley Gutierrez, Akiko Shimamura, Daniel E. Bauer

## Abstract

Despite extensive investigation, the molecular control of developmental hemoglobin expression remains incompletely elucidated. Hemoglobin switching is controlled by transcription factors, miRNAs, and RNA-binding proteins (RBPs) that enforce gene regulatory changes through development. Here we examine the role of the heterochronically silenced N-6 methyladenosine (m6A) RNA-binding protein IGF2BP1 that was previously described to regulate *HBG1/2* indirectly by suppressing *BCL11A* expression through an unknown mechanism. We find that IGF2BP1 binds and activates HIC2, itself a BCL11A repressor. Furthermore, we identify that IGF2BP1 plays a BCL11A-independent role by direct binding to *HBG1/2* to promote its translation. Stop codon-proximal m6A-modified coding sequences within *HBG2* transcripts are necessary and sufficient for direct positive regulation mediated by IGF2BP1. This work deepens the mechanistic understanding of hemoglobin switching and suggests a physical relationship between heterochronic RBPs and globin transcripts.

## Introduction

The developmental timing of globin gene expression and, of particular clinical interest, the switch from fetal to adult hemoglobin, is a coordinately controlled process, whereby the fetal *HBG1/2* genes are silenced and adult *HBB* gene is reciprocally activated around birth^1–3^. This transition is mediated by genes which are themselves heterochronically regulated and support not just the fetal to adult hemoglobin switch but developmentally regulated gene expression programs in many cell contexts^4–10^. These heterochronic factors function in a network regulating and regulated by *IGF2BP1* and include the transcription factor *HIC2* (directly regulating *BCL11A* transcription), *HMGA2* (unclear mechanism), *LIN28B* (regulating *BCL11A* translation), and adult-stage *let-7* miRNA (regulating *HIC2, LIN28B, IGF2BP1* and *HMGA2*)^4–8,10,11^. *HIC2* was identified as a regulator of HbF expression that represses *BCL11A* transcription^4^. Additionally, the fetal factor *LIN28B* was shown to suppress *BCL11A* translation^5^. *HMGA2* overexpression leads to increased HbF in adult erythroblasts through an unclear mechanism. Notably, the miRNA *let-7* targets all of these heterochronic HbF regulators including *IGF2BP1*^5,7,11–14^.

The m6A RNA-binding protein *IGF2BP1* is developmentally silenced after the fetal stage across many tissues, including in erythroid precursors during the hemoglobin switch^8,15^. m6A modifications are written on RNAs by the *METTL3/METTL14* heteroduplex at DRACH motifs^16^. Most m6A that is deposited on transcripts is observed in the vicinity of the stop codon and in the 3’ UTR. Study of regulatory m6A has been mainly focused on the 3’ UTR^17–19^. Recent work has identified CDS-m6A decay, in which the m6A RNA-binding protein *YTHDF2* binds to stop codon-proximal m6A in the coding region to degrade transcripts^20^. Before the appreciation of m6A as a dynamic regulatory mark and the identification of IGF2BP1 as an m6A RNA-binding protein, the coding region determinant of instability (CRD) in *MYC* mRNA was characterized as a target of *IGF2BP1* (initially known as CRD-BP), which binds to and stabilizes the otherwise unstable MYC transcript, suggesting that *IGF2BP1* in addition to *YTHDF2* can contribute to control of gene expression through CDS m6A^21–24^.

BCL11A is a key orchestrator of hemoglobin switching, binding to the *HBG1/2* promoters, repressing *HBG1/2* transcription^25–28^. Mutations in the binding sites of BCL11A at the -115 region of the *HBG1/2* promoters have been found to be causative of HPFH^28^. BCL11A recruits the NuRD complex to the *HBG1/2* gene loci to enable globin switching^33^. An erythroid-specific enhancer controlling *BCL11A* gene expression is the target of the first FDA-approved CRISPR gene therapy for transfusion-dependent β-thalassemia (TDT) and sickle cell disease (SCD)^29–32^.

Although m6A was first identified as a post-transcriptional RNA modification in 1972, the identification of FTO as an m6A demethylase decades later led to the understanding that m6A is a dynamically regulated mark with a diverse set of roles involved in post-transcriptional control of gene expression^35–37^. The m6A writing complex is a heteroduplex composed of METTL3 and METTL14, aided by the cofactor WTAP, which interacts with other cofactors^37^. FTO and ALKBH5 have been identified as m6A demethylases^37^. In addition to writers and erasers of m6A, readers include IGF2BP1/2/3, YTHDF1/2/3, YTHDC1/2, H2RNPC and HNRNPA2B1^19,37–39^. Binding of IGF2BP family RBPs is generally associated with increased translation and/or RNA stability, whereas binding of YTHDF family proteins is generally associated with destabilization and RNA decay; however, the gene regulatory outcome for individual transcripts or contexts may not be predictable or completely understood^40,41^. Notably, loss of expression of the *METTL3* gene component of the m6A writer complex leads to hematopoietic defects in both zebrafish and mice^23^.

Mechanistically, IGF2BP1 supports the expression of bound mRNAs by increasing their mRNA stability and/or translation efficiency^40^. In erythroid cells, however, *IGF2BP1* overexpression has been shown to have a role in depressing the expression of the key globin switching transcription factor *BCL11A* in adult erythroid cells. IGF2BP1 was shown to bind to *BCL11A* mRNA yet *BCL11A* mRNA stability and translation were unchanged, suggesting an uncertain mechanism^8,29,42,43^. Also, in this previous report, IGF2BP1 overexpression resulted in HbF levels up to 90% of total hemoglobin, notably higher than previously demonstrated upon loss of *BCL11A* expression^8,44^. Both the downregulation of BCL11A, opposite to the typical outcome of *IGF2BP1* binding to a target mRNA, as well as the magnitude of the HbF upregulation upon *IGF2BP1* overexpression raise the question of additional mechanisms besides loss of *BCL11A* expression.

## Results

### *BCL11A-*independent regulation of fetal hemoglobin by IGF2BP1

To investigate the role of IGF2BP1 in adult stage globin expression, we overexpressed *IGF2BP1* in immortalized human umbilical-derived erythroid progenitors (HUDEP) cells with an adult erythroid gene expression profile (HUDEP-2)^45^. We observed an increase in *HBG1/2* mRNA level, consistent with previous findings (**Figure 1A**). To test whether *BCL11A* was required for this upregulation in fetal hemoglobin expression, we overexpressed IGF2BP1 in *BCL11A* knockout (Δ*BCL11A*) HUDEP-2 cells. These cells have increased baseline *HBG1/2* mRNA and HbF levels as expected. Nonetheless, upon IGF2BP1 overexpression, we observed a further increase in *HBG1/2* mRNA and of HbF levels as compared to empty vector control cells (**Figure 1B and C**). This suggests a role for *IGF2BP1* in regulating HbF expression that is *BCL11A*-independent in addition to the previously described regulation of *BCL11A* by *IGF2BP1*.

**Figure 1.**
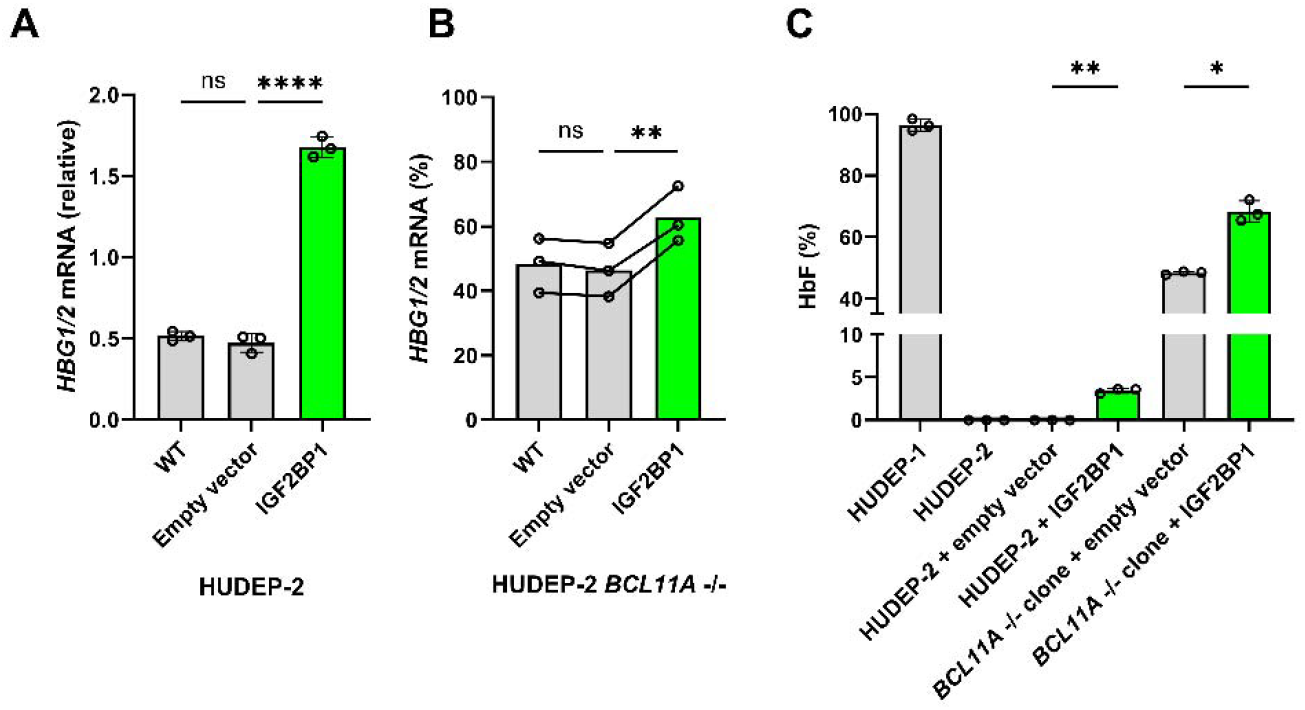
BCL11A-independent regulation of fetal hemoglobin by IGF2BP1. **A:** Expansion phase HUDEP-2 cells were transduced with control or IGF2BP1 expression vector using lentivirus. Primers targeting HBG1/2 and HBB were used to measure the RNA level by RT-qPCR vs. catalase. **B:** Expansion phase HUDEP-2 BCL11A-/- clones were transduced and RT-qPCR was performed. HBG1/2 mRNA level is reported as a percent of total HBB+HBG1/2 mRNA. **C:** Indicated cell lines were erythroid differentiated in vitro for 12 days and hemoglobin HPLC was performed to quantify the level of HbF expressed as percentage of HbF+HbA. All p values were calculated using one-way ANOVA. *p < 0.05; **p < 0.01; ***p < 0.001; ****p < 0.0001; ns, not significant.

### *IGF2BP1* overexpression indirectly regulates *BCL11A* by supporting *HIC2* expression

m6A is a dynamically regulated RNA modification for which most described regulatory sites for protein coding transcripts are found within the 3’ UTR, and most observed m6A modifications lie within the 3’ UTR or in proximity to the stop codon in the last coding exon^18,36,37,46–48^. The regulatory outcome of any given m6A mark is primarily driven by the identity of the m6A RNA-binding protein that binds the mark^37,38^. IGF2BP1 is an m6A RNA-binding protein which supports the RNA stability and/or the translation efficiency of bound targets^14,19,40,49,50^. *IGF2BP1* overexpression suppresses *BCL11A* expression, which raises the possibility that *IGF2BP1* may indirectly control *BCL11A* expression^8^.

Recent work has identified a role for the transcription factor HIC2 in repression of *BCL11A* transcription, and *HIC2* is itself silenced by the heterochronic adult-stage miRNA *let-7*, which is also known to negatively regulate embryonic/fetal-stage factors *LIN28B* and *IGF2BP1*^4,12,51^. We speculated that *HIC2* may be the direct target of *IGF2BP1* responsible for the *IGF2BP1*-mediated suppression of *BCL11A* upon *IGF2BP1* overexpression in adult erythroblasts. *IGF2BP1* overexpression in HUDEP-2 cells led to an upregulation of HIC2 protein as well as a reduction in BCL11A protein (**Figure 2**). To test whether the reduction in *BCL11A* expression upon *IGF2BP1* overexpression required *HIC2*, we generated *HIC2* -/-HUDEP-2 clones. Overexpression of *IGF2BP1* in these cells showed blunted reduction of BCL11A expression, suggesting that *HIC2* is required for the effect of *IGF2BP1* overexpression on BCL11A (**Figure 2B**). Overexpression of *IGF2BP1* in HUDEP-2 cells led to increased *HIC2* expression as measured by RT-qPCR (**Figure 2C**). To determine whether *IGF2BP1* binds directly to *HIC2* mRNA, we performed IGF2BP1 CLIP-qPCR on HUDEP-2 cells overexpressing *IGF2BP1* and observed enrichment of *HIC2* mRNA (**Figure 2D**). *HIC2* appears to be the primary effector whereby IGF2BP1 mediates *BCL11A* silencing.

**Figure 2.**
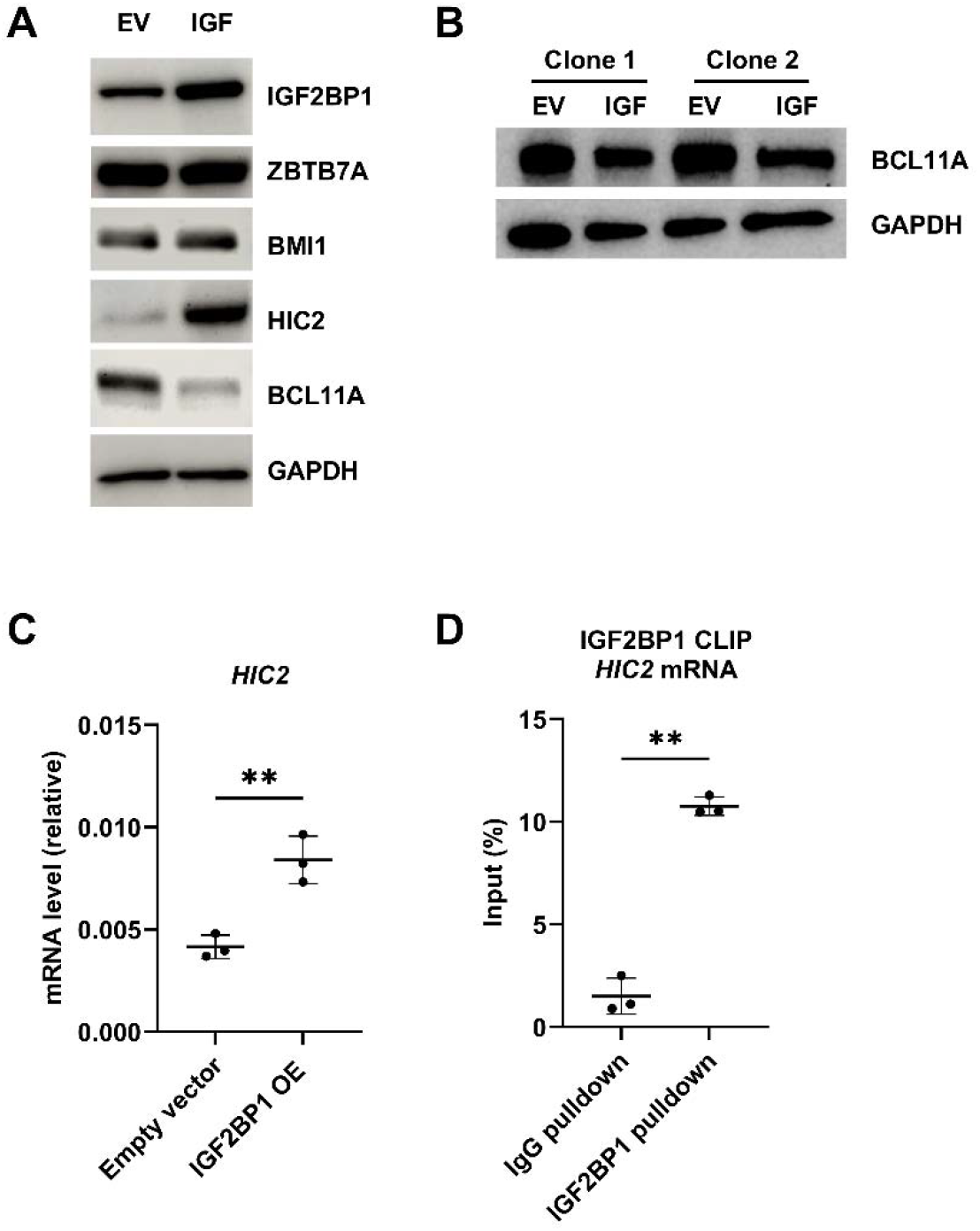
The BCL11A-dependent effect of IGF2BP1 on HbF regulation is mediated by HIC2. **A:** Expansion phase HUDEP-2 cells were transduced with control or IGF2BP1 overexpression vectors and protein was isolated for western blotting of indicated proteins. **B:** HUDEP-2 HBG2-2A-mCherry ΔHIC2 cells were transduced with control or IGF2BP1 overexpression and protein was isolated for western blotting of indicated proteins. **C:** HIC2 mRNA level from HUDEP-2 cells transduced with control or IGF2BP1 overexpression vector was measured by RT-PCR relative to catalase. **D:** IGF2BP1 binding of HIC2 mRNA level was measured by IGF2BP1 CLIP-qPCR.

### *IGF2BP1* directly regulates *HBG1/2 mRNA* post-transcriptionally

To identify candidate *IGF2BP1* target genes responsible for the *BCL11A*-independent regulation of HbF, we performed differential gene expression analysis upon overexpression of *IGF2BP1* in HUDEP-2 cells. Additionally, we compared these datasets to RNA-seq data from primary human fetal liver, cord blood, adult bone marrow as well as wild-type HUDEP-1 and HUDEP-2 cells. Overexpression of *IGF2BP1* resulted in a shift of HUDEP-2 cells towards a fetal-like expression profile primarily driven by increases in the expression of developmentally regulated hemoglobin gene expression, specifically *HBZ, HBE1, HBG1/2*, and the *HBBP1* pseudogene (**Figure S1**). Profiling of known fetal hemoglobin regulators through this method failed to reveal obvious candidates responsible for the *BCL11A*-independent regulation. Given its function as a positively regulating RBP, we hypothesized that IGF2BP1 might directly bind to *HBG1/2*. We performed *IGF2BP1* CLIP-qPCR from HUDEP-1 cells, HUDEP cells with a fetal-like expression pattern (**Figure 3A**)^45^. We observed significant binding of IGF2BP1 to its known target *HMGA1* (**Figure 3B**). We also found significant binding of *IGF2BP1* to the *HBG1/2* mRNA when measured using primers for an amplicon spanning the exon 2/exon 3 junction of *HBG1/2* (**Figure 3C**) as compared with the negative control histone mRNA *H2AB1* (**Figure 3D**). A separate pair of primers targeting *HBG1/2* exon 3 CDS yielded a similar result in terms of IGF2BP1 binding (**Figure 3E**). These results raised the possibility of a direct role for *IGF2BP1* in regulating *HBG1/2* post-transcriptionally.

**Figure 3.**
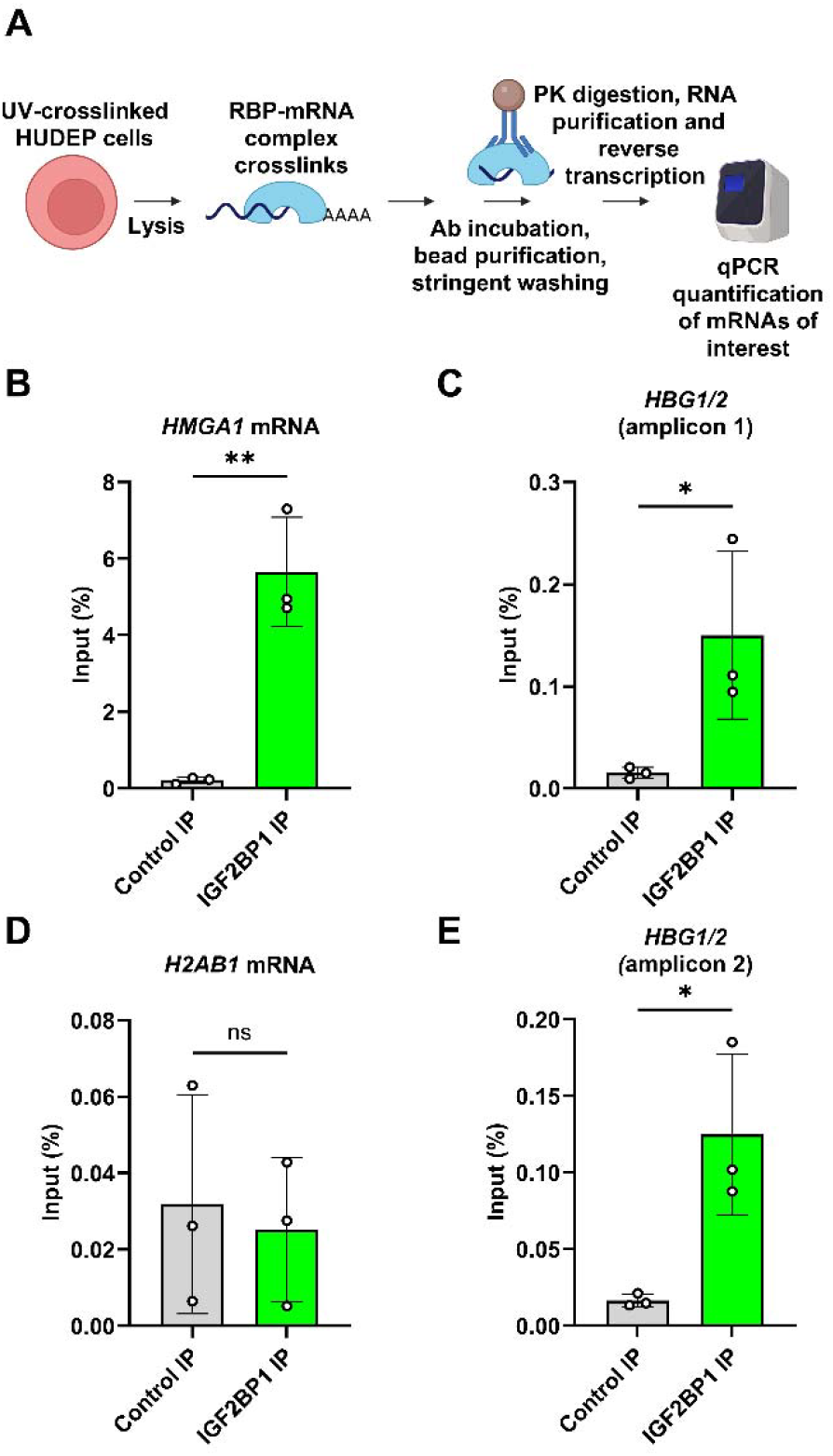
IGF2BP1 CLIP-qPCR. **A:** Schematic of the IGF2BP1 CLIP-qPCR experiment with expansion phase HUDEP-1 cells. **B:** IGF2BP1 CLIP-qPCR measuring HBG1/2 with primers spanning the exon 2/exon 3 junction, **C:** HBG1/2 exon 3 targeting primers, **D:** known IGF2BP1 target HMGA1 and **E:** H2AB1 mRNA were used to assess IGF2BP1 binding. Control IPs are performed with isotype-matched antibody.

### *IGF2BP1* regulates *HBG1/2* mRNA directly through a stop codon-proximal DRACH motif

To distinguish the indirect *BCL11A*-dependent effect of *IGF2BP1* on *HBG1/2* transcription from the possible direct regulatory effects of *IGF2BP1* binding to the *HBG1/2* mRNA, we designed a lentiviral reporter system independent of endogenous globin gene transcription. This *SFFV-HiBiT-HBG-PGK-eGFP* reporter construct employs a HiBiT tag at the 5’ end of the *HBG2* protein coding region which enables transgene-specific mRNA and protein level quantification. To test whether IGF2BP1 can upregulate *HiBiT-HBG* expression, we transduced HUDEP-2 cells with the *HiBiT-HBG* reporter vector (**Figure 4A**). Then the culture was split before subsequent transduction with IGF2BP1 or control vector. Upon IGF2BP1 overexpression, we observed an increase in luminescence as compared with control cells (**Figure 4B**). To test whether this effect was *BCL11A* independent, we transduced *BCL11A -/-* HUDEP-2 clones. We observed a similar increase in HiBiT-HBG luminescence upon *IGF2BP1* overexpression in *BCL11A -/-* HUDEP-2 clones, demonstrating that this regulation is *BCL11A* independent (**Figure 4C**). This effect size was similar to the experiment in *BCL11A* wild-type cells. Next, we tested whether this regulation was intact in primary CD34+ hematopoietic stem and progenitor cells (HSPCs). We transduced CD34+ HSPCs from 3 donors with the *HiBiT-HBG* reporter vector and control or *IGF2BP1* overexpression vector and observed a similar increase in HiBiT-HBG luminescence upon IGF2BP1 overexpression throughout *in vitro* erythroid differentiation (**Figure 4D**). These results suggest that *HBG1/2* regulation by IGF2BP1 is independent from the endogenous transcriptional regulation by *BCL11A* or other transcriptional regulators of the endogenous β*-*globin gene cluster.

**Figure 4.**
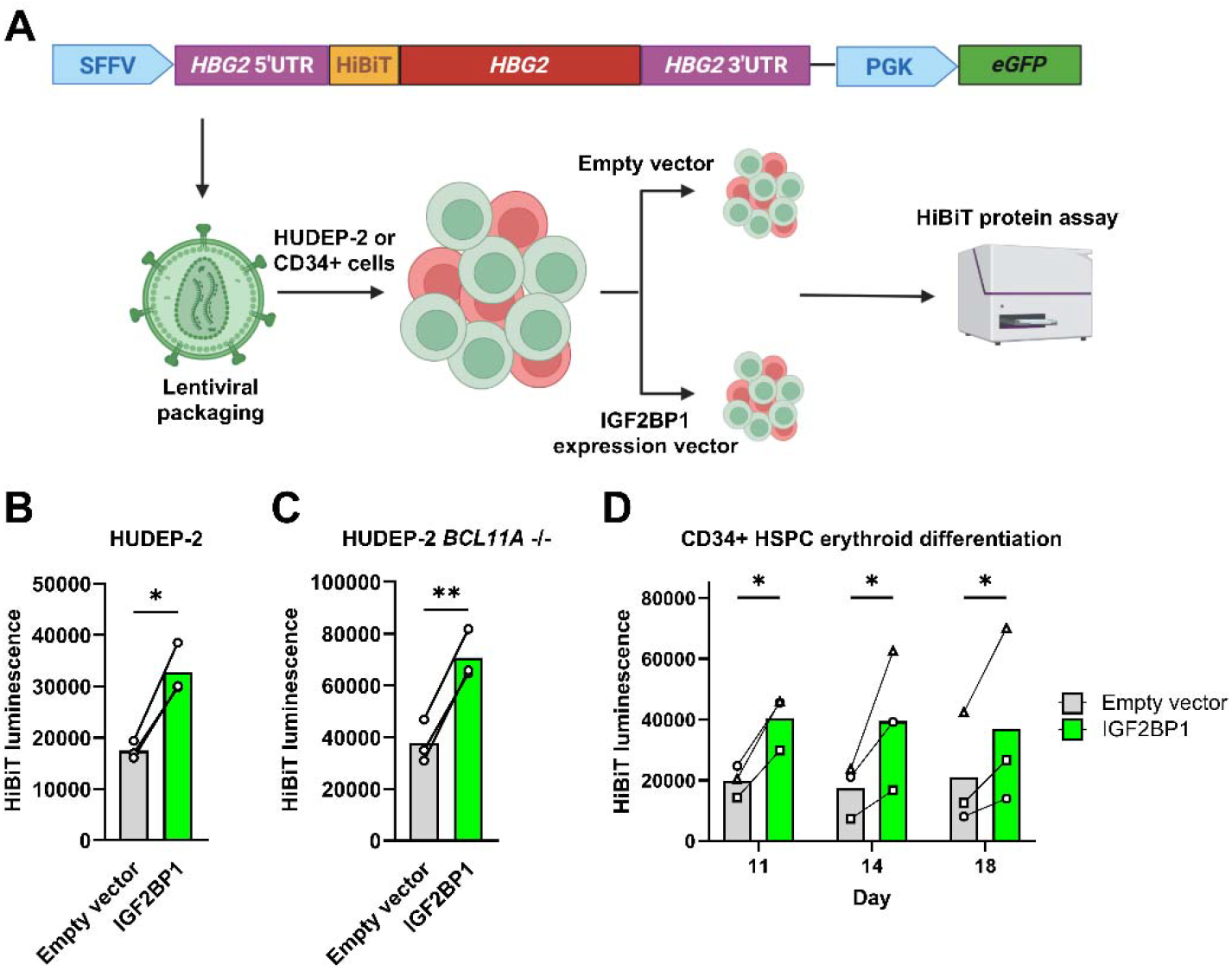
IGF2BP1 regulates HBG1/2 level independent of endogenous transcriptional regulation. **A:** Lentivirus containing the SFFV-HiBiT-HBG-PGK-eGFP cassette was used to transduce HUDEP-2 or CD34+ HSPCs. Each transduction was split for a second round of transduction with IGF2BP1 or empty control vector, then luminescence was measured following selection for **B:** Wild-type HUDEP-2 cells, **C:** ΔBCL11A HUDEP-2 cells, and **D:** in vitro erythroid differentiated CD34+ HSPCs measured at the indicated timepoints.

IGF2BP1 is an m6A-RNA binding protein that acts to stabilize its targets and promote their expression by improving translation efficiency^19,21,40,52,53^. Given the observations that IGF2BP1 directly binds to and promotes the expression of *HBG1/2* mRNA, we performed polysome profiling to test whether this *BCL11A*-independent mechanism acted by promoting translation. HUDEP-2 cells were transduced with *SFFV-HiBiT-HBG2-PGK-eGFP* reporter lentivirus (*HiBiT-HBG)* before further transduction with IGF2BP1 or empty vector. Lysates from these cells were subjected to polysome profiling and were fractionated into twenty-four fractions (**Figure 5B**). Following fractionation, we isolated RNA and measured the abundance of *HiBiT-HBG* reporter mRNA in each fraction. We grouped fractions corresponding to the following three groups: polysome free, early polysome, and late polysome. We observed a significant increase in the abundance of reporter mRNA in late polysome fractions upon IGF2BP1 overexpression consistent with increased translation of the transgenic *HiBiT-HBG* mRNA mediated by IGF2BP1 (**Figure 5**).

**Figure 5.**
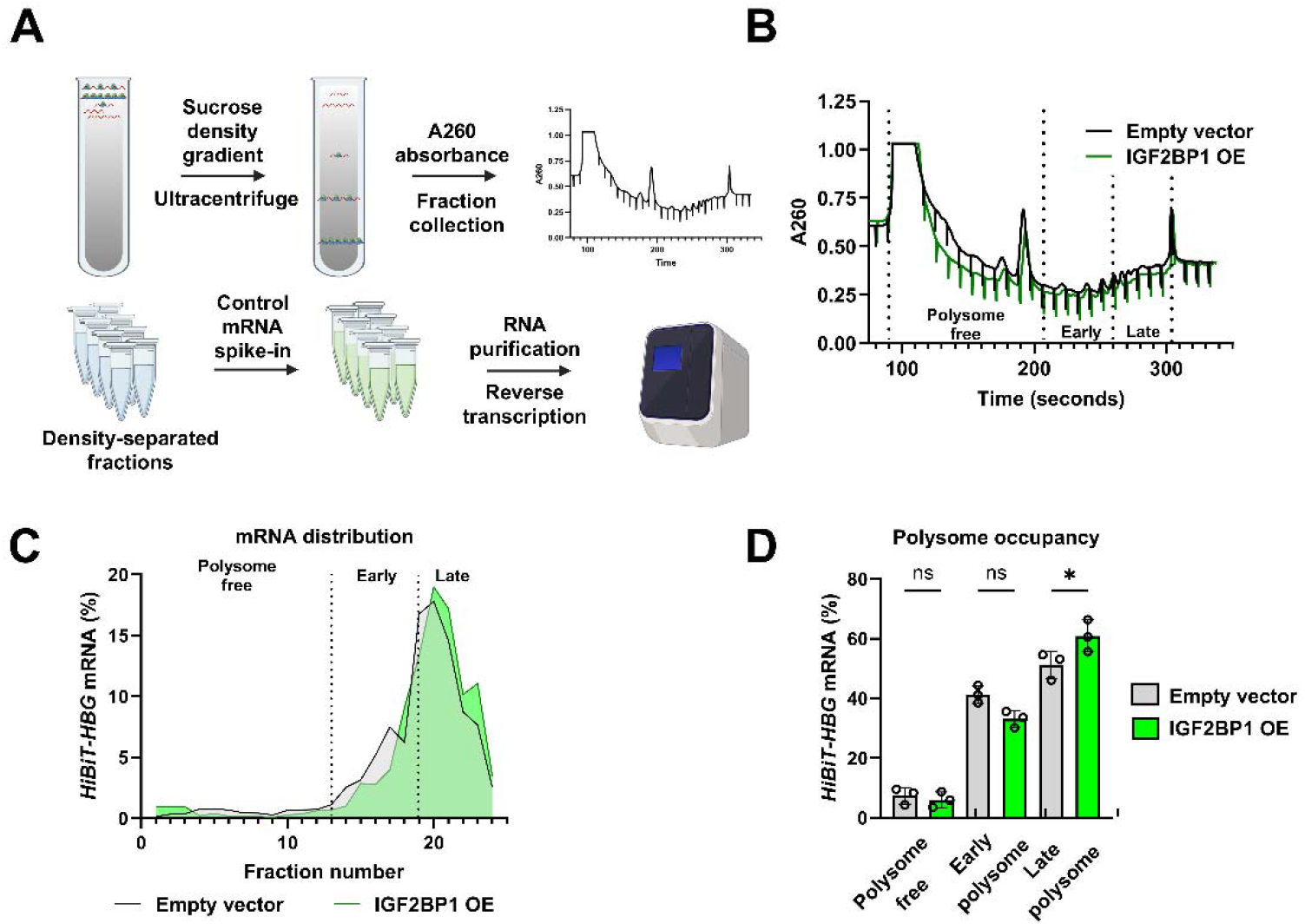
IGF2BP1 promotes translation of HBG1/2 mRNA. Expansion phase HUDEP-2 cells transduced with HiBiT-HBG reporter virus were split and further transduced with IGF2BP1 or control overexpression virus and selected with puromycin. **A:** Polysome profiling was performed using sucrose gradient ultracentrifugation and fractionated into 24 fractions to separate highly translated (more dense) translation complexes. **B:** Overlay of representative polysome profiles. Polysome free, early polysome, and late polysome are indicated. **C:** Distribution of the HiBiT-HBG mRNA across the polysome profile. Dashed lines separate the polysome free, early polysome, and late polysome regions of the profile. **D:** Indicated mRNA content in the polysome free, early polysome, and late polysome regions.

Given the described function of *IGF2BP1* as an m6A RNA-binding protein that supports the translation of bound transcripts, we searched for motifs matching that of the *METTL3/METTL14* m6A writer complex motif DRACH in the *HBG1/2* mRNAs. To our surprise, no m6A sites are present in the 3’ UTR sequences of *HBG1* or *HBG2* where functional m6A elements have typically been described across the transcriptome. We observed 10 putative m6A motifs in *HBG1/2*, each located in shared coding sequences (CDS) (**Figure 6**).

**Figure 6.**
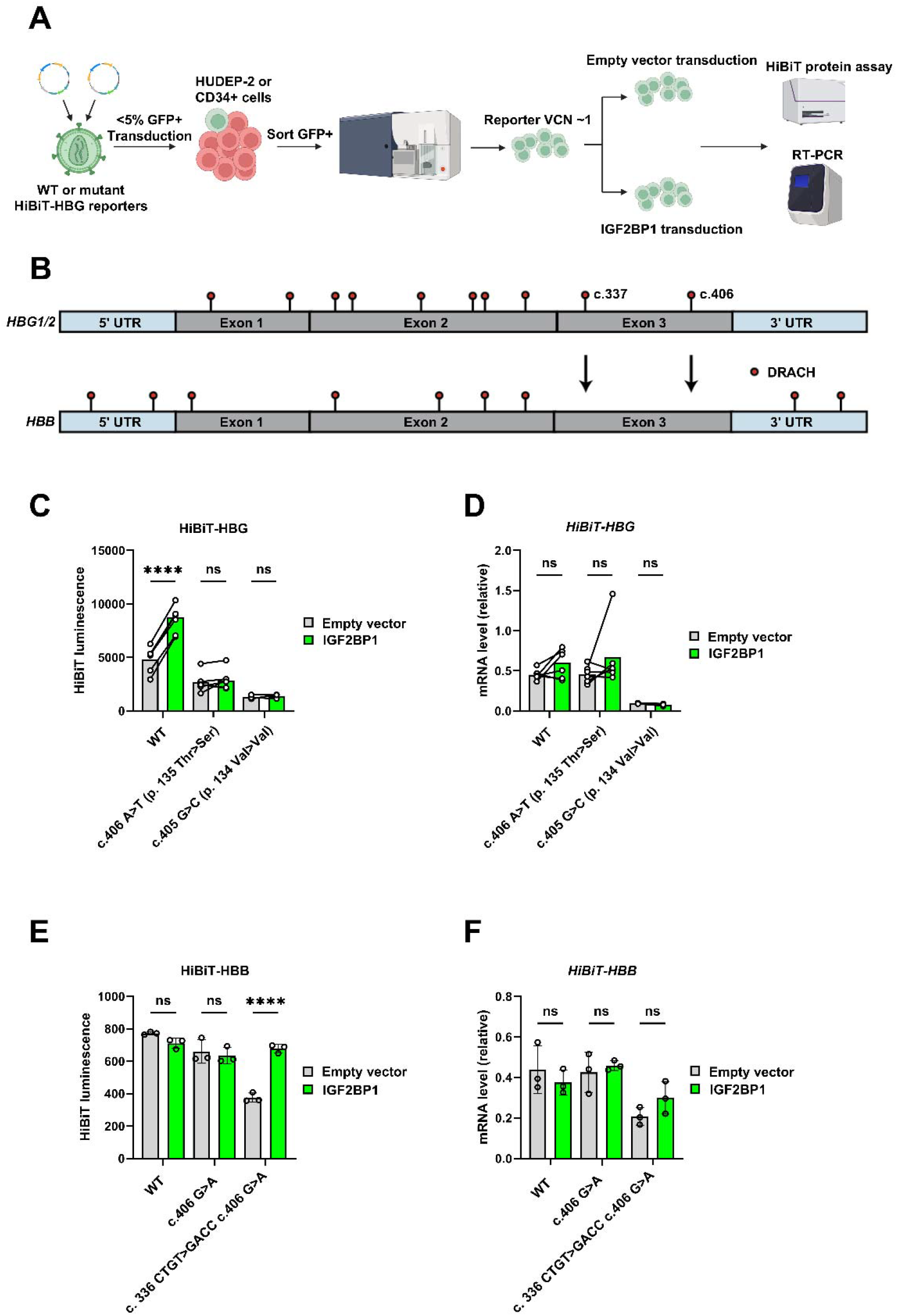
IGF2BP1 promotes expression of HBG1/2 through the stop codon-proximal DRACH motif. **A:** Lentivirus carrying the wild-type or stop codon-proximal DRACH mutant SFFV-HiBiT-HBG-PGK-eGFP cassette was used to transduce wild-type HUDEP-2 cells at low efficiency to ensure low vector copy number to enable comparisons between different variants. GFP+ cells were sorted and transduced with either IGF2BP1 or control overexpression vector and selected with puromycin. **B:** An alignment schematic of HBG1/2 and HBB mRNAs with DRACH motifs indicated by lollipops. The position of the stop codon-proximal m6A motifs refers to the position of the central ‘A’ in the DRACH motif. **C:** HiBiT-HBG protein level was measured by HiBiT luminescence. Wild-type but not stop codon-proximal mutants were responsive to IGF2BP1 overexpression. **D:** HiBiT-HBG mRNA levels were measured relative to GAPDH in cells at timepoints matched to **C. E:** Wild-type, c.406 G>A (p.135 A>T) variant, or c. 336 CTGT>GACC (p.112 C>T) c. 406 G>A (p.135A>T) HiBiT-HBB cells were generated as described in **A**. HiBiT-HBB protein level was measured by HiBiT-luminescence. **F:** HiBiT-HBB mRNA levels were measured relative to GAPDH in cells at timepoints matched to **E**.

We hypothesized that putative m6A motifs in exon 3 CDS could provide selective RNA binding of *HBG1/2*, given that they are stop codon-proximal and absent in *HBB* despite the high degree of homology between these transcripts^18,40^. Of these, we selected the most stop codon-proximal putative m6A motif and designed *HiBiT-HBG* reporter vectors containing the nonsynonymous mutation c.406A>T (DRACH>DRTCH, p.135Thr>Ser), eliminating the possibility of m6A at this position by replacing the target adenine in the motif, or c.405 G>C (DRACH>DCACH, p.134Val>Val), a synonymous mutation disrupting the second position in the m6A motif.

Given our observation that IGF2BP1 bound to *HBG1/2* transcripts (**Figure 3**), we investigated whether the introduction of point mutations at the stop codon-proximal m6A motif could disrupt or mitigate the binding of IGF2BP1 to the *HiBiT-HBG* reporter transgene. CLIP-qPCR in HUDEP-2 cells overexpressing IGF2BP1 with the c.406A>T mutant reporter demonstrated a reduction in IGF2BP1-bound c.406A>T *HiBiT-HBG* reporter transcript compared to WT reporter transcript (**Figure S2**).

To test the impact of these point mutations on gene expression, we transduced HUDEP-2 cells at low multiplicity (<5% transduction frequency) to ensure that different constructs all had comparable vector copy number of ∼1. We observed that only the WT HiBiT-HBG could but not the c.406A>T or c.405G>C mutants could be upregulated by IGF2BP1 overexpression (**Figure 6**). Notably, the baseline protein level of both mutants was lower than the WT reporter vector. We also tested the mRNA level of the HiBiT reporter transcript from time-matched samples upon IGF2BP1 overexpression. We observed that IGF2BP1 failed to increase the transcript level of any of the three reporter constructs consistent with a role for IGF2BP1 in directly supporting the translation of *HBG1/2* through its stop codon-proximal m6A motif.

### Addition of HBG-like stop codon-proximal m6A motifs to *HBB* enables post-transcriptional regulation by *IGF2BP1*

We observed that the *HBB* transcript lacks a homologous stop codon-proximal m6A motif as the one we mutated in *HBG1/2* to disrupt *IGF2BP1-*mediated post-transcriptional regulation. Notably, it differs at the paralogous globin position by only a single base-pair. The naturally occurring Calvino *HBB* variant (c.406G>A, p.135Ala>Thr) is a hematologically silent *HBB* variant which creates a ‘*HBG1/2*-like’ m6A motif at this position^54^. We created a *WT* and *c*.*406G>A* version of a *HiBiT-HBB* reporter. However, we observed no change in its mRNA or protein level in response to *IGF2BP1* overexpression (**Figure 6E-F**). Some targets of IGF2BP1 and other m6A reader proteins bind at multiple methylated sites on target mRNA^21,40^. Notably, *HBG1/2* has 2 putative m6A motifs in exon 3 of its CDS that *HBB* lacks (**Figure 6B**). We next tested whether addition of a second gamma-like m6A motif to the *HBB* reporter could enable IGF2BP1-mediated post-transcriptional regulation, termed the *c*.*336CTGT>GACC, c*.*406G>A HiBiT-HBB* reporter. Upon IGF2BP1 overexpression, the *c*.*336CTGT>GACC, c*.*406G>A HiBiT-HBB* reporter, unlike the *WT* or *c*.*406G>A* reporters, exhibited a similar upregulation response to the *WT HiBiT-HBG* reporter at the protein level while the mRNA level remained unchanged (**Figures 6E-F**). These results suggest that these two stop codon proximal CDS m6A sites are sufficient to confer IGF2BP1-mediated post-transcriptional regulation.

## Discussion

As part of hemoglobin switching, PRC1 acts to transcriptionally silence at least 3 heterochronic components supporting the fetal gene expression program – *IGF2BP1, IGF2BP3*, and *LIN28B*^6^. In 2017, *IGF2BP1* was first identified as a regulator of HbF, found to be associated with reduced BCL11A levels. Several lines of evidence suggest that its repression of BCL11A may be indirect, including an effect size exceeding that typically observed following *BCL11A* loss, a lack of change in *BCL11A* translation or RNA stability upon IGF2BP1 overexpression, and the typical role of *IGF2BP1* in supporting the expression of its targets^8,15^. Our current work demonstrates that *IGF2BP1* acts through a bipartite mechanism to regulate HbF expression. We show that the component of this regulation dependent upon *BCL11A* occurs via control of *HIC2*. Recent work has shown that *HIC2* is a target of *let-7* miRNA, and that *HIC2* acts as a direct transcriptional repressor of *BCL11A*^4,51^. *IGF2BP1* has been shown to inhibit miRNA-mediated inhibition of mRNA targets. Our work identifies a role for *IGF2BP1* in supporting the expression of *HIC2*, thus indirectly repressing *BCL11A* transcription. When *IGF2BP1* is introduced to adult erythroid cells where it is typically silenced, this leads to reactivation of HbF expression.

The *BCL11A*-independent mechanism of IGF2BP1 acting directly on *HBG1/2* mRNA identified in this work is supported by observations of IGF2BP1 binding, translation regulation, and the necessity and sufficiency of stop-codon proximal m6A motifs. Notably, *HBB* lacks the m6A motifs present in *HBG1/2*. While m6A modification has become appreciated as a dynamic regulator of gene expression, most well-understood examples have been characterized within the 3’ UTR^37,48,55^. Our report adds to an emerging body of work pointing towards CDS m6A as a regulatory feature, in this instance a mechanism supporting the expression of *HBG1/2* mRNA driven by m6A marks in the CDS bound by the m6A reader *IGF2BP1*. Several lines of evidence support the importance of CDS m6A, particularly in the vicinity of the stop codon. Firstly, the conserved nature of some marks; secondly, several well characterized examples from RNA biology before the explosion of work on m6A in the 2010s (particularly *c-MYC*, one of the first well characterized targets of *IGF2BP1*); and thirdly, recent studies characterizing a novel developmentally associated RNA decay mechanism termed CDS-m6A decay (CMD), which requires the m6A reader YTHDF2^18,20,22,52,53,56^.

We speculate that *IGF2BP1* binding may help to suppress CMD by interfering with YTHDF2 binding to the developmentally regulated *HBG1/2* mRNA, a protective measure unneeded by *HBB*, which lacks stop codon-proximal m6A sites in the CDS that may promote translational pausing. This would be consistent with a similar mechanism proposed for *IGF2BP1* target cMYC prior to the current understanding of m6A as a dynamic regulatory mark - specifically, that *IGF2BP1* binding (also known as CRDBP) may bind to the ∼250 nucleotide coding region determinant of instability (CRD), would protect a downstream endonucleolytic cleavage site within the CRD that could become exposed by ribosomal pausing (thought at that time to be due to rare codon usage within the element) ^20,21^.

In conclusion, we have identified that the reintroduction of *IGF2BP1* in adult erythroid cells supports the expression of the developmentally regulated *HBG1/2* genes through a bipartite mechanism. Firstly, there is an indirect transcriptional regulation whereby *IGF2BP1* supports *HIC2* expression directly enabling the transcriptional silencing of *BCL11A*. Notably, *HIC2* mRNA is a target of let-7, and *IGF2BP1* has been reported to have a role in inhibiting miRNA-mediated regulation by binding near miRNA binding sites in the 3’ UTR of transcripts^50,51^. Second, *IGF2BP1* directly binds to *HBG1/2* transcripts at stop codon-proximal m6A motifs, and this binding is required to support their translation. Introducing *HBG*-like m6A motifs to the *HBB* transcript boosts its IGF2BP1-dependent expression. In addition to our observation of IGF2BP1-mediated direct post-transcriptional support of *HBG1*/2 expression, recent work has identified the RNA-binding protein *PUM1*, a *KLF1* target gene, as a negative post-transcriptional regulator of *HBG1*^57^. These results suggest a previously underappreciated layer of hemoglobin switching regulation, that is, a physical relationship between heterochronic RBPs and developmentally regulated globin genes.

## Methods

### HUDEP cell culture

*Expansion phase culture:* HUDEP-1 and HUDEP-2 cells and derivative cell lines were cultured in expansion phase media and split 1:10 approximately every 3 days. Since HbF levels may be sensitive to cell culture stress, cells were maintained at a concentration below 1e6 cells/mL. HUDEP cells require doxycycline to maintain expression of the E6/E7 system driven by the tetracycline-inducible promoter^45,58^.

*Differentiation phase culture:* HUDEP-1 and HUDEP-2 cells and derivative cell lines were cultured first in expansion phase media, then transferred to EDMII for 8 days. At the end of 8 days, cells were transferred to EDMIII for up to 4 additional days of culture.

*Expansion phase media* – StemSpan Serum-Free Expansion Media (SFEM, Stemcell Technologies) supplemented with stem cell factor (SCF, 50 ng/mL), erythropoietin (EPO, 3 IU/mL), doxycycline (DOX, 1 mcg/mL), dexamethasone (DEX, 0.4 mcg/mL) and penicillin/streptomycin (Pen/Strep, Gibco, 2% final concentration).

*Erythroid Differentiation Media II* (EDMII) – Iscove’s Modified Dulbecco Media plus 1% L-glutamine and 2% Pen/Strep (IMDM, Life Technologies) supplemented with holo-human transferrin (330 mcg/mL), recombinant human insulin (10 mcg/mL), heparin (2 IU/mL), inactivated human plasma (5%), EPO (3 IU/mL), SCF (100 ng/mL), DOX (1 mcg/mL).

*Erythroid Differentiation Media III* (EDMIII) *–* IMDM plus 1% L-glutamine and 2% Pen/Strep supplemented with holo-human transferrin (330 mcg/mL), recombinant human insulin (10 mcg/mL), heparin (2 IU/mL), inactivated human plasma (5%), EPO (3 IU/mL), DOX (1 mcg/mL).

### CD34+ HSPC in vitro erythroid differentiation

CD34+ HSPCs from healthy donors were obtained from the Fred Hutch Cancer Center. CD34+ HSPCs were differentiated in a 3-phase culture system consisting of culture from days 0-7 in EDMI followed by EDMII from days 7-11 and concluding with culture in EDMIII from days 11-18. CD34+ HSPCs were treated with 8uM cyclosporine H to enable more efficient lentiviral transduction^59^.

*CD34+ Erythroid Differentiation Media III* (EDMIII) *–* IMDM plus 1% L-glutamine and 2% Pen/Strep supplemented with holo-human transferrin (330 mcg/mL), recombinant human insulin (10 mcg/mL), heparin (2 IU/mL), inactivated human plasma (5%), and EPO (3IU/mL).

*CD34+ Erythroid Differentiation Media I* (EDMI) *–* CD34+ EDMI is composed of CD34+ EDMIII supplemented with hydrocortisone (1 mcM), SCF (100 ng/mL), and IL-3 (5 ng/mL).

*CD34+ Erythroid Differentiation Media II* (EDMII) *–* CD34+ EDMII is composed of CD34+ EDMIII supplemented with SCF (100 ng/mL).

### Hemoglobin HPLC

Hemoglobin HPLC was performed on indicated cells following RBC lysis using the HBA_2_/F assay on the Biorad D-10 Dual assay system. HbF percentages are expressed as a percent of HbF to HbA (HbF/(HbF+HbA)).

### Western blot

Samples were collected and lysed in RIPA lysis buffer supplemented with cOmplete protease inhibitor cocktail (Millipore Sigma), then quantitated with Pierce BCA protein assay kit according to the manufacturer’s instructions. 10 mcg of protein was loaded per sample for SDS-PAGE gel electrophoresis prior to transfer of protein to PVDF membranes using Bio-Rad Trans-Blot Turbo Transfer system. Antibodies used in this study can be found in the supplemental materials.

### RT-qPCR

RNA was isolated using RNeasy mini kit (Qiagen) purification or Trizol extraction according to manufacturer recommendations. Reverse transcription was performed using iScript (Biorad) which contains a mix of random hexamers and oligo-dT as the primer for first strand synthesis. SYBR select master mix (ThermoFisher) was used with a final primer concentration of 250 nM to prepare the reactions for cycling and measurement using the Quantstudio 3 Real-Time PCR system (Applied Biosystem). Primer pairs are listed in the supplemental material.

### IGF2BP1 CLIP-qPCR

2e7 suspension cells were collected and washed 2X in PBS, resuspended in 10 mL of PBS, plated, and crosslinked at 4000 mcJ/cm^2^, pelleted, and flash frozen. Within 2 weeks, cells were lysed in fresh iCLIP lysis buffer (50 mM Tris-HCl pH 7.4, 100 mM NaCl, 1% NP-40, 0.1% SDS, 0.5% sodium deoxycholate (protect from light), 1:200 Protease Inhibitor Cocktail III (Pierce) supplemented with RNase inhibitor. Lysates were split in half for matched pulldown with IGF2BP1 or control antibody pulldown. Dynabeads M280 Sheep anti-mouse beads were equilibrated with iCLIP lysis buffer and coupled to antibodies before mixing with cell lysates and were incubated overnight. Following incubation, samples were washed alternating between high salt (50 mM Tris-HCl pH 7.4, 1M sodium chloride, 1 mM EDTA, 1% NP-40, 0.1% SDS, 0.5% sodium deoxycholate and low salt (20 mM Tris-HCl 7.4, 10 mM MgCl2, 0.2% Tween-20) wash buffers for a total of 8 washes. RNA was liberated from the beads by proteinase K digestion followed by heat inactivation, then purified using RNA clean and concentrator columns (Zymo) alongside matched input samples, then measured by RT-qPCR. Protein was isolated by treatment of beads with 2X Laemmli buffer and analyzed by western blot.

### Polysome profiling

HUDEP-2 cells transduced with *PLVX-IGF2BP1-Puro* or control empty vector were treated with cycloheximide at a concentration of 100ug/mL to arrest translation elongation for 10 minutes at 37 degrees before cell lysis at 2e7 cells per 1mL of polysome lysis buffer (20 mM HEPES pH 7.4, 100 mM KCl, 10 mM MgCl2, 1% NP40, 1% DOC, 1X Complete Mini EDTA Free, 1mM Pefabloc, 1.4 uM Pepstatin A, 100 ug/ml cycloheximide, 200 ug/ml Heparin, 2 ul/ml RNAsin. 125 mcL of clarified lysate was loaded onto 10-50% sucrose gradients and ultracentrifuged in an SW-55 Ti rotor at 45k RPM prior to fractionation. Firefly luciferase mRNA prepared by *in vitro* transcription (NEB T7 HiScribe) was spiked into each fraction prior to RNA purification by trizol purification, reverse transcription (iScript, Biorad) and RT-PCR.

### Lentiviral production

293T cells cultured in 15 cm plates were transfected using linear PEI MW ∼25,000 (Kyfora) with lentiviral packaging and envelope vectors and a lentiviral expression plasmid. After 2 days, supernatant containing lentivirus was collected and concentrated by ultracentrifugation at 24,000 RPM for 2 hours in an SW28 rotor through a 20% sucrose cushion and resuspended in OPTI-MEM.

### HiBiT luminescence assays

The *HiBiT-HBG2* and *HiBiT-HBB* versions of the *SFFV-HiBiT-PGK-eGFP* were cloned into the PLVX backbone from synthetic gene fragments that included the 5’ and 3’ UTR from each gene respectively (Genewiz), as well as the HiBiT peptide tag at the 5’ end of the CDS (Promega). Fragments were joined by Gibson assembly using NEBuilder HiFi DNA assembly mix (NEB)^59^. Variants of these vectors were cloned using site-directed mutagenesis with mismatch primers to obtain variants of interest. HUDEP cells or CD34+ HSPCs transduced with HiBiT-globin lentivirus were collected for analysis using the Nano-Glo HiBiT Lytic Detection System (Promega) according to the manufacturer’s instructions. Except for low VCN experiments, cells were first transduced with globin vector before being subsequently transduced by control or IGF2BP1 overexpression vector after 2 days to control for HiBiT-globin transduction level. For low VCN experiments, cells were transduced with HiBiT-globin lentivirus such that fewer than 10% of cells were GFP positive, then sorted for GFP positivity using BD FACS Aria and subsequently transduced with control or IGF2BP1 overexpression vector and selected with puromycin. 50k cells per well were analyzed in opaque white 96-well microplates and luminescence was measured using the Clariostar Plus microplate reader. Each biological replicate was measured in technical triplicate.

### RNAseq

HUDEP-1, HUDEP-2, transduced with *PLVX-SFFV-IGF2BP1-Puro* or empty vector were differentiated as described in EDMII for 8 days, then collected in Trizol for RNA purification before Truseq library construction. Libraries were sequenced on an Illumina Novaseq partial lane paired-end 150 bp sequencing run performed with Novogene. For analysis of this data and of primary cell RNAseq data, adapters were trimmed with Cutadapt, pseudomapped and counted with Salmon, and differential gene expression analysis was performed with DESeq2^60,61^. This dataset is available on NCBI’s Gene Expression Omnibus (GEO) through GEO Series accession number GSE303945.

## Acknowledgements

S.C. was supported by NIDDK (F31DK122637) and NHLBI (T32HL066987). D.E.B. was supported by the Doris Duke Foundation (#2022092), the St. Jude Children’s Research Hospital Collaborative Research Consortium, and the National Institutes of Health (R01HL167513). HSPCs were obtained from Fred Hutch Cooperative Center of Excellence in Hematology (U54DK106829). We thank Partha Das, Rajesh Gunage, Chad Harris, Pim Rullens, Anna Deck, Stuart Orkin, Richard Gregory, and John Manis for helpful discussions and assistance and the HSCI-BCH Flow Cytometry Research Lab for technical support.

## Author contributions

Conceptualization, S.C. and D.E.B.; Methodology, S.C. and D.E.B.; Investigation, S.C., T.W., G.H., M.H., D.V., M.S., J.Z., F.A., A.G., A.S., and D.E.B., Original Draft, S.C., Writing, Review, and Editing, S.C. and D.E.B., Funding Acquisition, D.E.B., Supervision, A.S. and D.E.B.

## Supplemental information

**Figure S1.**
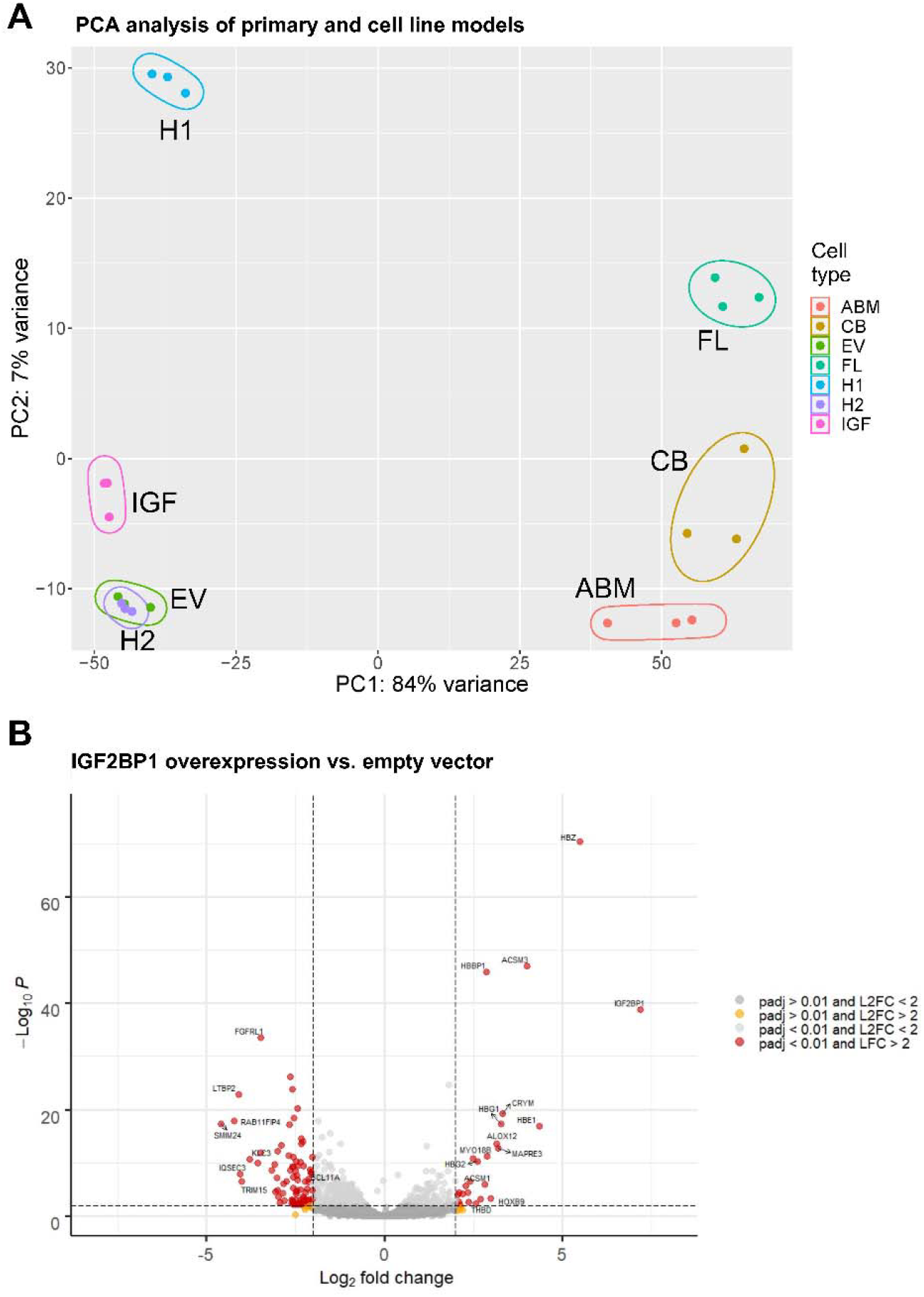
IGF2BP1 promotes a fetal-like expression profile. HUDEP-2 cells were transduced with control or IGF2BP1 overexpression lentiviral vectors. RNA samples were prepared from HUDEP-1 (H1), HUDEP-2 (H2), HUDEP-2 + empty vector (EV), and HUDEP-2 + IGF2BP1 overexpression (IGF) cells and used for RNAseq, then compared with pre-existing RNAseq data from adult bone marrow (ABM), cord blood (CB) and fetal liver (FL) derived erythroid precursors. A: Principal component analysis of HUDEP and primary cell RNAseq. B: Volcano plot of HUDEP-2 + empty vector and HUDEP-2 + IGF2BP1 OE.

**Figure S2.**
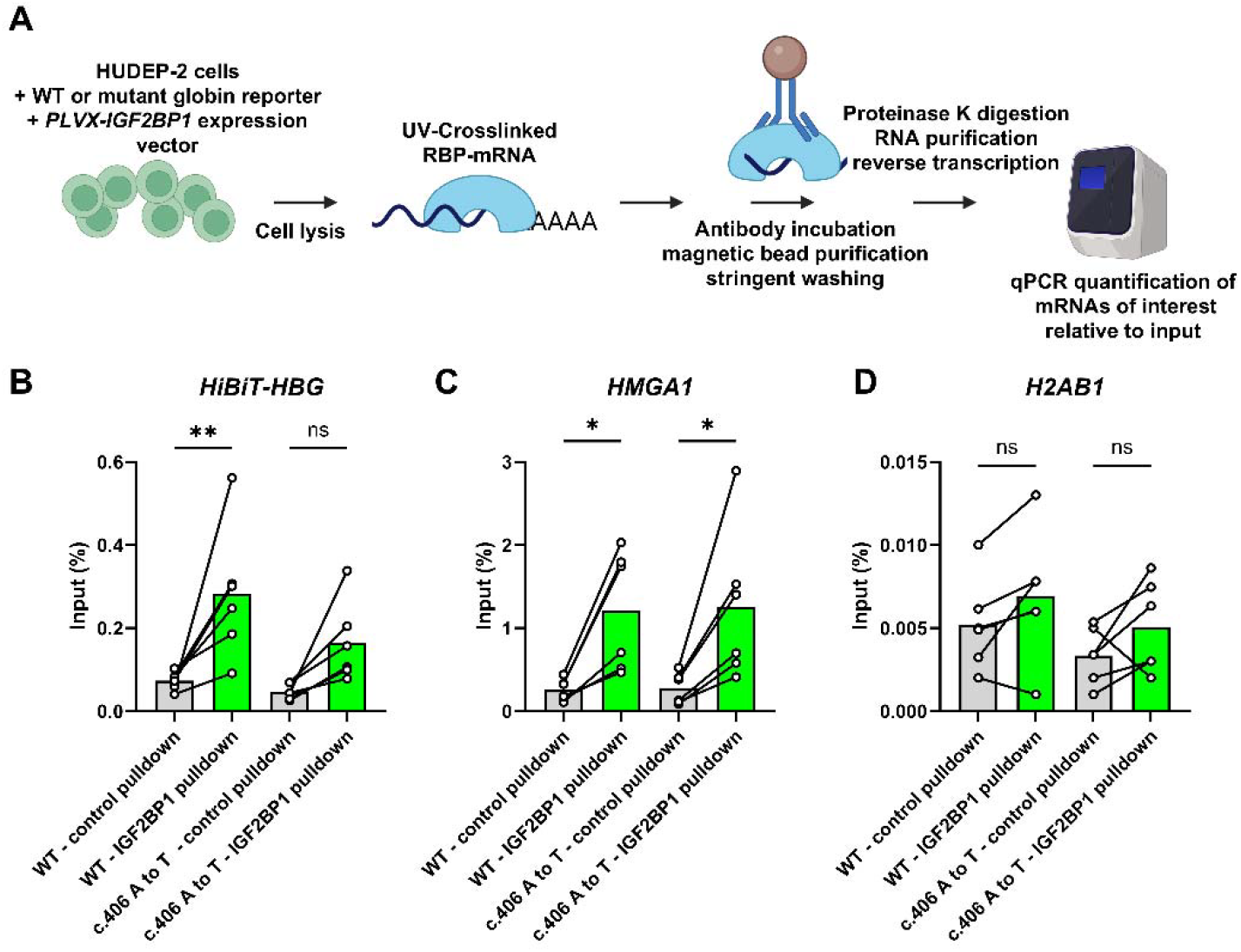
Mutagenesis of the stop codon-proximal DRACH motif in HBG1/2 mRNA reduces IGF2BP1 binding. **A:** HUDEP-2 cells with wild-type or c.406 A to T mutant HiBiT-HBG reporter vector overexpressing IGF2BP1 were used to perform IGF2BP1 CLIP-qPCR. Enrichment of **B:** HiBiT-HBG mRNA **C:** known IGF2BP1 target HMGA1 **D:** negative control H2AB1 mRNA.

### Primers

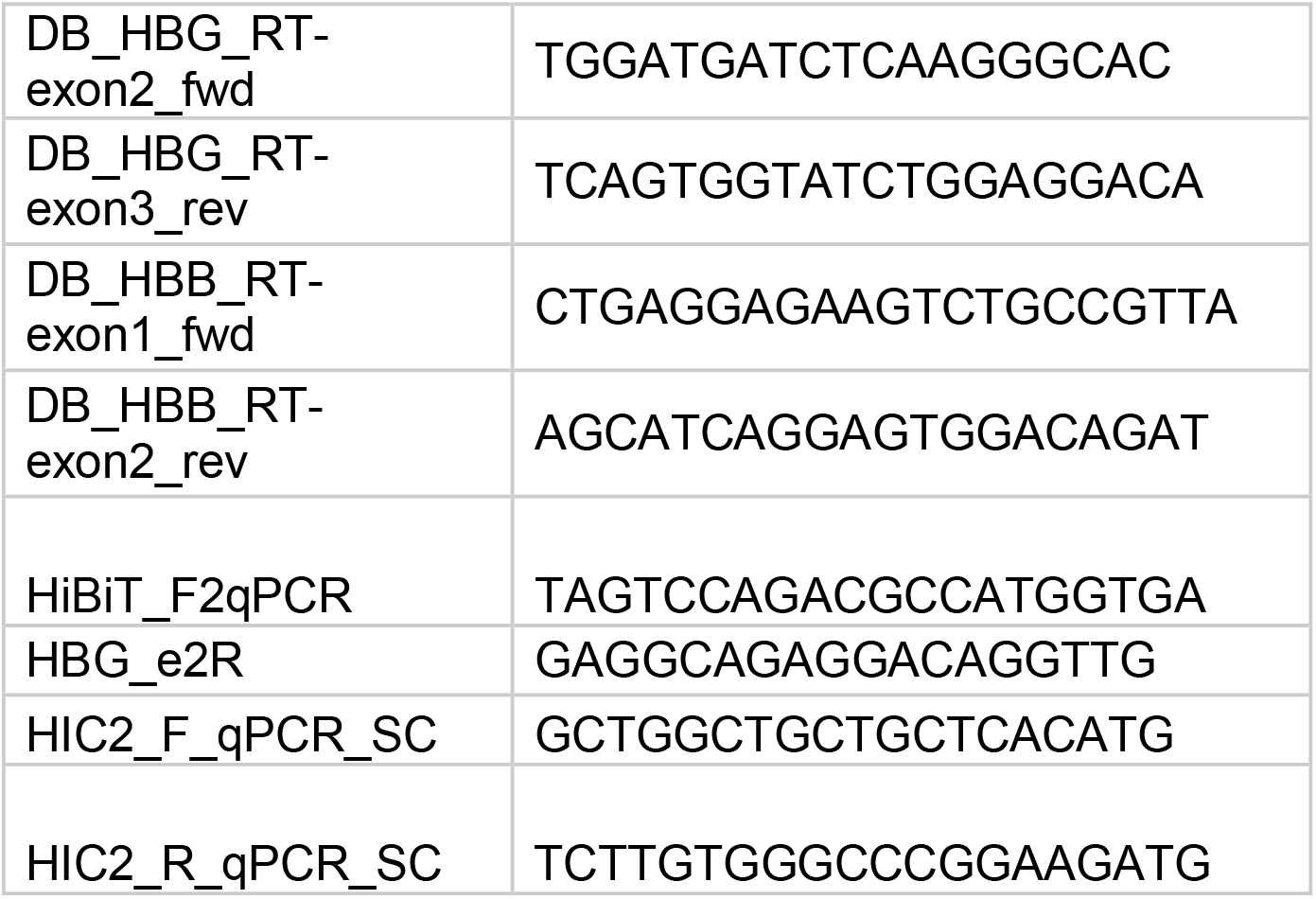
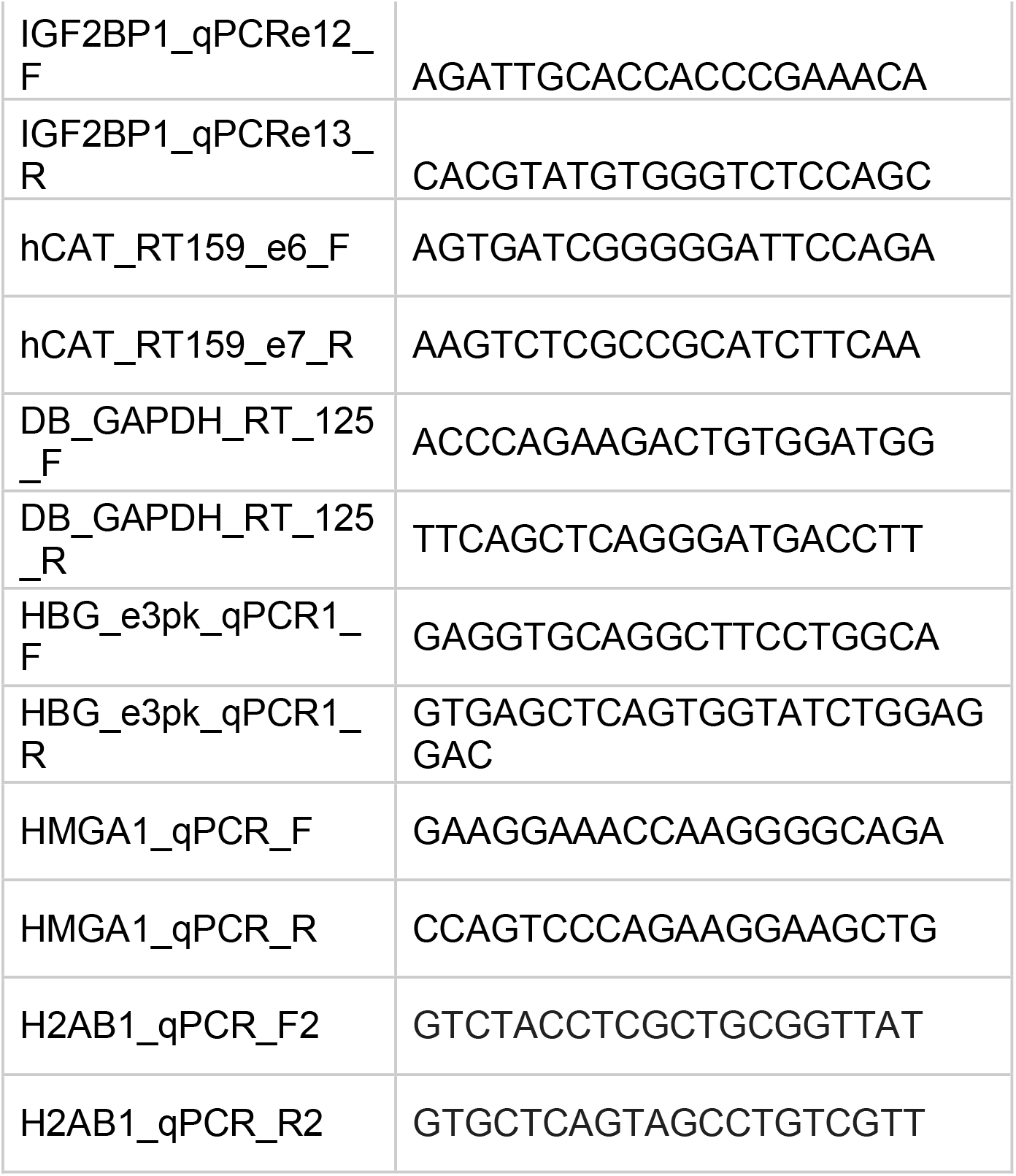

### Antibodies

|  |  |  |
| --- | --- | --- |
| Cell signaling technology | 8482S | IMP1 antibody |
| Abcam | ab191401 | Recombinant Anti-Ctip1/BCL-11A antibody [EPR14943-44] |
| Sigma | 14-3309-82 | Pokemon (LRF) Monoclonal Antibody (13E9) |
| CST | #5856S | BMI1 antibody |
| Cell Signaling Tech | 5174S | GAPDH (D16H11) XP® Rabbit mAb |
| Proteintech | 22788-1-AP | HIC2 Polyclonal antibody |

